# Designer *Sinorhizobium meliloti* strains and multi-functional vectors for direct inter-kingdom transfer of high G+C content DNA

**DOI:** 10.1101/449991

**Authors:** Stephanie L. Brumwell, Michael R. MacLeod, Tony Huang, Ryan Cochrane, Rebecca S. Meaney, Maryam Zamani, Ola Matysiakiewicz, Preetam Janakirama, David R. Edgell, Trevor C. Charles, Turlough M. Finan, Bogumil J. Karas

## Abstract

Storage and manipulation of large DNA fragments is crucial for synthetic biology applications, yet DNA with high G+C content can be unstable in many host organisms. Here, we report the development of *Sinorhizobium meliloti* as a new universal host that can store DNA, including high G+C content, and mobilize DNA to *Escherichia coli, Saccharomyces cerevisiae*, and the eukaryotic microalgae *Phaeodactylum tricornutum*. We deleted the *S. meliloti hsdR* restriction-system to enable DNA transformation with up to 1.4 x 10^5^ efficiency. Multi-host and multi-functional shuttle vectors (MHS) were constructed and shown to stably replicate in *S. meliloti, E. coli, S. cerevisiae*, and *P. tricornutum*, with a copy-number inducible *E. coli* origin for isolating plasmid DNA. Crucially, we demonstrated that *S. meliloti* can act as a universal conjugative donor for MHS plasmids with a cargo of at least 62 kb of G+C rich DNA derived from *Deinococcus radiodurans*.

## Introduction

The field of synthetic biology aims to utilize existing or novel biological parts and systems to create organisms that can help address global problems including the increased demand for food, fuel, therapeutics, and high value chemicals. However, one of a major obstacles in synthetic biology is that many organisms of interest lack genetic tools such as autonomously replicating plasmids, well characterized promoters/terminators, selective markers, genome-editing tools, and protocols to uptake and install DNA [1,2]. This problem can be addressed by cloning whole chromosomes or large DNA fragments of a donor genome in a surrogate host, where genetic tools are in place and manipulations can be performed [3,4]. Currently, the most common host for the capture and manipulation of DNA fragments is *Saccharomyces cerevisiae*, however returning cloned or engineered fragments to destination cells is still challenging due to the lack of direct transfer methods from *S. cerevisiae*, such as bacterial conjugation. In addition, *S. cerevisiae* cannot maintain large DNA fragments with G+C content >40% without additional engineering [5–7], and many industrially useful bacterial strains have a G+C content above this range. For example, *Streptomyces* species have a G+C content >65% and are important for the production of antibiotics (gentamicin, kanamycin, tetracycline, etc.) [8]. Therefore, it is desirable to develop a host to clone, maintain, manipulate, and transfer large DNA fragments, including high G+C content, to bacterial and eukaryotic host cells.

We selected *Sinorhizobium meliloti* as a host due to its high G+C content genome (62%), and available origins of replication that could be used to maintain large DNA fragments [9]. *S. meliloti* model strain Rm1021 has a multipartite genome including a chromosome (3.65 Mb), pSymA megaplasmid (1.35 Mb), and pSymB chromid (1.68 Mb) [10]. Recently, a derivative of *S. meliloti* with a minimal genome lacking the pSymA and pSymB replicons was developed, resulting in a 45% reduction of the genome [11]. Two essential genes were identified in the pSymB chromid, *engA* and tRNA^arg^, and these genes were transferred to the main chromosome [12]. Multiple derivatives of *S. meliloti* now exist that vary in genome size, nutrient requirements and generation time [11]. The replication origins of pSymA and pSymB were identified and characterized [13], and can be used to generate designer replicating plasmids. Additionally, several genetic tools have been developed for this species including vectors based on *repABC* origins taken from various α-Proteobacteria that can be maintained and selected for in *S. meliloti* [14]. Three of these vectors can be maintained within wildtype *S. meliloti* at one time, along with the endogenous pSymA and pSymB replicons [14]. Utilizing these vectors, an *in vivo* cloning method with Cre/lox-mediated translocation of large DNA fragments to the *repABC*-based vector was established [14]. Lastly, *S. meliloti* is a Gram-negative α-Proteobacterium that has a symbiotic relationship with legume plants and fixes nitrogen in root nodules, and therefore is of high importance in agriculture.

Here, we utilized the reduced genome strains of *S. meliloti* [11] to create designer strains with the restriction-system removed and the conjugative pTA-Mob [15] plasmid installed. pTA-Mob, which contains all of the genes needed for direct transfer via conjugation, can mobilize any plasmid containing an origin of transfer *(oriT).* Previous studies have demonstrated DNA transfer methods from *S. meliloti* through introduction of a Ti plasmid to plant species through mobilization of TDNA [16]. In addition, *S. meliloti* has been used to move DNA via conjugation in tri-parental matings [14].

In addition, we used identified origins from *S. meliloti* pSymA and pSymB replicons to create multi-host shuttle (MHS) vectors (named pAGE1.0, 2.0, 3.0, and pBGE1.0, 2.0, 3.0) that replicate in *S. meliloti, Escherichia coli* (by addition of a copy-number inducible bacterial artificial chromosome vector), *S. cerevisiae* (by addition of a yeast artificial chromosome vector) and *Phaeodactylum tricornutum* (by addition of a selective marker and yeast ARS). In addition, all plasmids contain an origin of transfer *(oriT)* that is acted on by proteins encoded on the conjugative plasmid, and is necessary for mobilization of the plasmid to the recipient cell [15,17]. The MHS vectors were tested for replication and stability in *S. meliloti.* Additionally, we demonstrated the ability to maintain large DNA fragments on these plasmids, using pAGE2.0 (18.5 kb) to clone a complete 62 kb native plasmid from *Deinococcus radiodurans* (G+C content 57%) [18]. These plasmids were moved into *S. meliloti* strains via an optimized electroporation protocol and via a newly developed polyethylene glycol (PEG) method. Above all, we have developed protocols to directly transfer the MHS plasmids via conjugation from *S. meliloti* to *E. coli, S. cerevisiae*, and *P. tricornutum.*

## Results and Discussion

### Development of designer bacterial strains

To develop *S. meliloti* as a host we used the ΔpSymA strain that retained the pSymB chromid, as this strain had the fastest doubling time when compared to the other reduced-genome strains [11]. First, the strains were engineered to remove the restriction-system (Δ*hsdR*) to allow for development of more efficient transformation methods. Disruption or deletion of the *hsdR* gene has been previously reported in the Rm1021 strain *of S. meliloti*, and deletion mutants were shown to have enhanced transformation efficiencies [19,20]. In our reduced strains, the *hsdR* gene was replaced by a FRT-Km/Nm-FRT cassette and the resulting Δ*hsdR*::Nm mutant allele was transferred to various background strains via transduction of Nmr. The FRT-Km/Nm-FRT cassette was then removed by introducing an unstable plasmid (pTH2505) carrying the Flp recombinase, followed by curing of this plasmid. The *hsdR* deletion strains were verified using diagnostic PCR with primers upstream and downstream of the *hsdR* gene (S1 Fig). *S. meliloti* RmP4098 ΔpSymA Δ*hsdR*, retaining the Km/Nm cassette, was transformed with the pTA-Mob plasmid [15] and used as the conjugative donor strain for all subsequent experiments (Fig 1a). In addition, reduced *S. meliloti* strains RmP3952 ΔpSymB Δ*hsdR* and RmP3909 ΔpSymAB Δ*hsdR* were developed using the same method described above.

**Fig 1.**
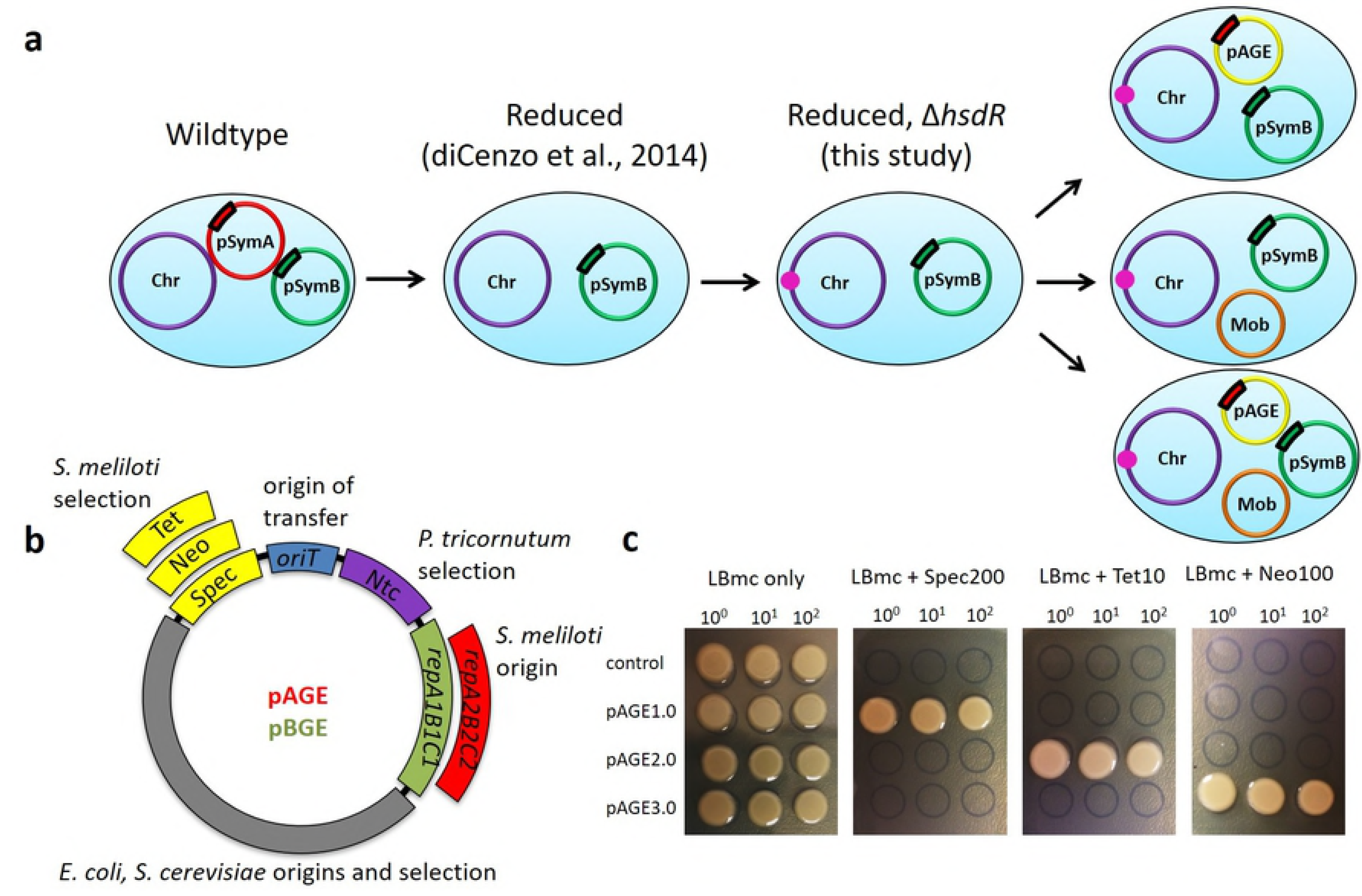
Development of designer *S. meliloti* strains and multi-functional vectors. (a) Schematic showing the creation of the designer *S. meliloti* ΔpSymA Δ*hsdR* strains including the genome reduction and deletion of the restriction-system. Several versions of the designer strains are available depending on the desired application including strains with the pTA-Mob conjugative plasmid, and/or compatible genome engineering plasmids (pAGE or pBGE) installed. (b) Plasmid map of pAGE/pBGE MHS vector with both standard and interchangeable components. Standard components include the pCC1BAC-yeast backbone, an origin of transfer *(oriT)*, and the selectable marker for *P. tricornutum* nourseothricin N-acetyl transferase (Ntc). Interchangeable components include selectable markers for *S. meliloti:* tetracycline (Tet), kanamycin/neomycin (Neo), and spectinomycin (Spec); and origins of replication for *S. meliloti:* the pSymA origin *(repA2B2C2)*, and pSymB origin *(repA1B1C1).* Three vectors (pAGE1.0, pAGE2.0, and pAGE3.0) were constructed utilizing the *repA2B2C2* origin with Spec, Tet or Neo selection, respectively. Three vectors (pBGE1.0, pBGE2.0, and pBGE3.0) were constructed utilizing the *repA1B1C1* origin with Spec, Tet or Neo selection, respectively. (c) Antibiotic selection test of three versions of pAGE plasmids in *S. meliloti* RmP4122 ΔpSymA Δ*hsdR* Nms. Antibiotic resistance includes Spec for pAGE1.0, Tet for pAGE2.0, and Neo for pAGE3.0.

Additionally, a designer *E. coli* strain was developed to simplify the current method of conjugation from *E. coli.* We used lambda red recombination to create an auxotrophic strain of *E. coli* Epi300, named ECGE101, by deleting the *dapA* gene. This gene is required for synthesis of diaminopimelic acid (DAP) [21], therefore ECGE101 requires DAP supplementation in the growth media and provides a useful, antibiotic-free method for counter-selecting *E. coli* after conjugation to *S. meliloti* or any other organism.

### Design, assembly and characterization of multi-host shuttle plasmids

With the reduced, restriction-system minus strains of *S. meliloti* created, and the origins of replications of the large megaplasmid and chromid (*repA1B1C1* and *repA2B2C2*) identified, multi-host shuttle (MHS) plasmids were developed. Six MHS plasmids were constructed to allow for stable replication and selection in *S. meliloti, E. coli, S. cerevisiae*, and *P. tricornutum*. These organisms were chosen as they are well-characterized model strains for bacterial, yeast and algal systems. These plasmids were constructed using a bacterial artificial chromosome (BAC) and yeast artificial chromosome (YAC) backbone [17]. The MHS plasmids contain an origin of replication captured from the native megaplasmid and chromid of *S. meliloti*, pSymA (*repA2B2C2*, added to pAGE vectors) or pSymB (*repA1B1C1*, added to pBGE vectors). The YAC also allows replication in *P. tricornutum* [17,22]. Selectable markers include spectinomycin (Sp), tetracycline (Tc), or kanamycin/neomycin (Km/Nm) resistance for *S. meliloti* (but also conferring some resistance in *E. coli*), nourseothricin N-acetyl transferase (Ntc) for *P. tricornutum*, HIS3 for *S. cerevisiae*, and chloramphenicol (Cm) resistance for *E. coli.* Finally, the plasmids contain an origin of transfer (*oriT*) that is necessary to allow for conjugation of these plasmids from *S. meliloti* to *E. coli, S. cerevisiae*, and *P. tricornutum* using the pTA-Mob helper plasmid (Fig 1b). Three versions of the pAGE vector differing only in the *S. meliloti* selective marker, pAGE1.0 (Sp), pAGE2.0 (Tc), and pAGE3.0 (Km/Nm), were constructed with the pSymA origin (*repA2B2C2*). An additional three pBGE vectors differing only in the *S. meliloti* selective marker, pBGE1.0 (Sp), pBGE2.0 (Tc), and pBGE3.0 (Km/Nm), were constructed with the pSymB origin (*repA1B1C1*). Once assembled [23], the plasmids were transformed into *E. coli* Epi300 (Epicentre) cells and a diagnostic digest was performed to confirm the correct assembly of the vectors (S2 Fig). The three pAGE plasmids were conjugated to *S. meliloti* RmP4122 ΔpSymA Δ*hsdR* Nms from *E. coli* ECGE101 conjugative strains, and transconjugants were spot plated on their respective antibiotic selections. We demonstrated plasmid replication of pAGE1.0, pAGE2.0, and pAGE3.0 in *S. meliloti* and their ability to provide resistance to Sp, Tc, and Nm, respectively (Fig 1c). Following transformation of pAGE2.0 into *S. meliloti* RmP4122 ΔpSymA Δ*hsdR* Nms, plasmid stability was tested by iterative subculturing every 10 generations to a total of about 50 generations. We observed, on average, that about 25% of plasmids were lost after 50 generations, as determined by the number of colonies unable to grow on selective media (LBmc 38 μM FeCl_3_ 2 μM CoCl_2_ Tc 5 μg/mL) after restreaking from nonselective media (LBmc 38 μM FeCl_3_ 2 μM CoCl_2_) (S3 Fig, S1 Table).

### Optimization of DNA transfer to *S. meliloti* via electroporation, a new polyethylene glycol transformation method and conjugation

In order to develop *S. meliloti* as a host, a highly efficient transformation method is required for the uptake of DNA. Currently, the most common transformation method used in *S. meliloti* is electroporation. Optimization of this method through transformation of *S. meliloti* RmP4122 ΔpSymA Δ*hsdR* Nms with three pAGE plasmids (∼18 kb) produced efficiencies averaging 1.4 x 10^5^ CFU μg^-1^ of DNA (Fig 2a). Additionally, since a PEG transformation method was successfully applied to move large DNA fragments (>1 Mb) in the transplantation protocol required to create the first synthetic cell [24], we developed a PEG transformation method in *S. meliloti* and were able to obtain efficiencies on average of 2.1 x 103 CFU μg^-1^ of DNA (Fig 2b, S4 Fig). In addition, conjugation has been previously established as a method of DNA transfer to *S. meliloti* [20,25], therefore we have developed an improved conjugation protocol from our conjugative *E. coli* ECGE101 strain carrying the pTA-Mob and pAGE1.0 plasmids. Using this method we obtained a conjugation efficiency averaging 4 transconjugants per every 10 recipient cells (S2 Table).

**Fig 2.**
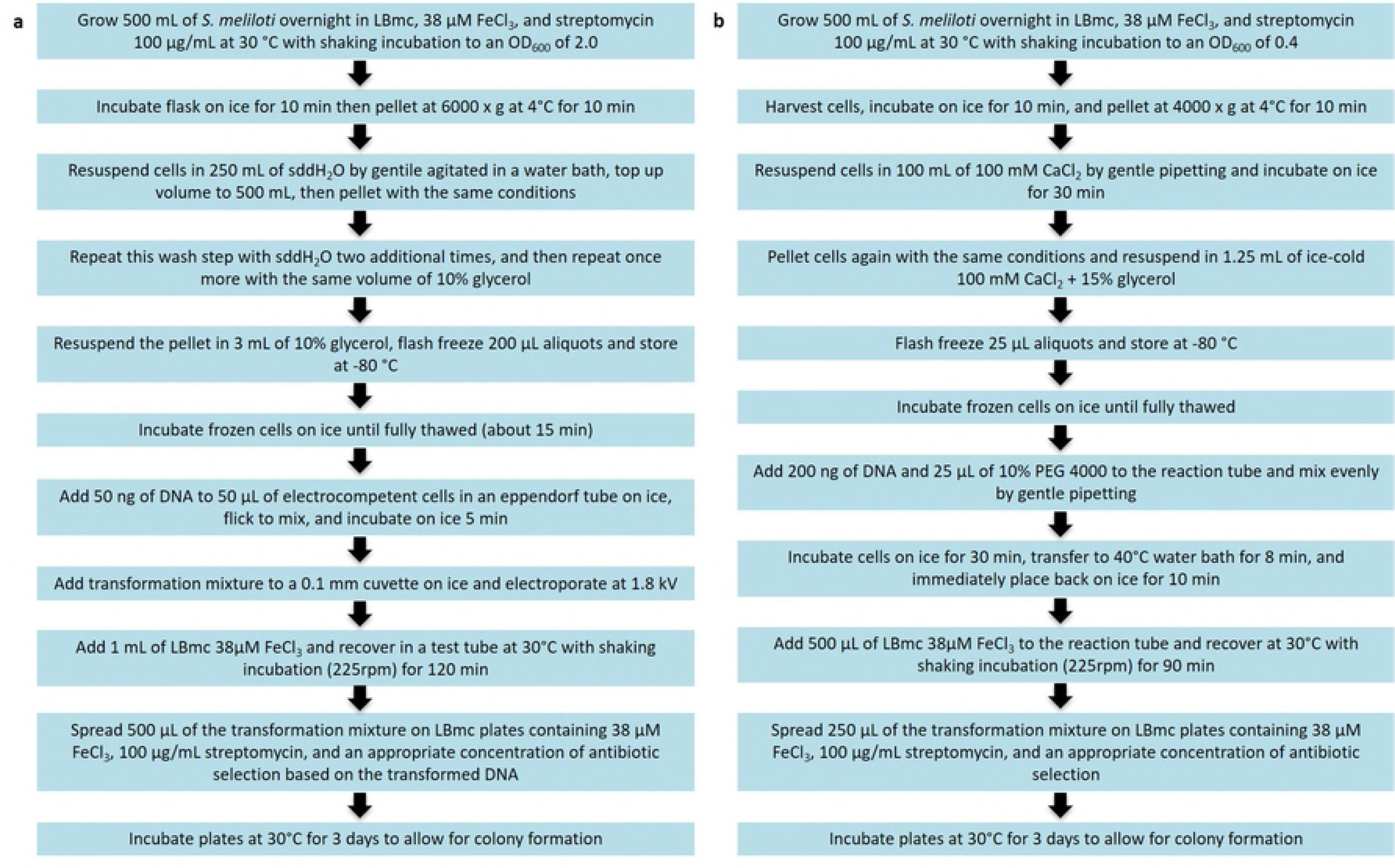
Workflow of optimized transformation protocols. (a) Preparation of competent cells for *S. meliloti* and electroporation protocol. (b) Preparation of competent cells for *S. meliloti* and PEG-mediated transformation protocol.

### Direct DNA transfer (via conjugation) from *S. meliloti* to *E. coli, S. cerevisiae* and *P. tricornutum*

We utilized the designer *S. meliloti* RmP4098 ApSymA *AhsdR* host strain carrying the pTA-Mob conjugative plasmid and pAGE1.0 compatible genome engineering plasmid to develop direct DNA transfer of pAGE1.0 (via conjugation) from *S. meliloti* to *E. coli, S. cerevisiae*, and *P. tricornutum* (Fig 3a-c,e). The ∼18 kb pAGE1.0 plasmid contains the pSymA origin *(repA2B2C2)* and the spectinomycin selectable marker for *S. meliloti.* Conjugation efficiencies from the *S. meliloti* conjugative donor to *E. coli, S. cerevisiae*, and *P. tricornutum* are 2.22 x 10^-1^, 7.99 x 10^-6^ and 9.40 x 10^-5^, respectively. After optimization, all conjugation experiments yielded a useable number of colonies (>20 per selective plate) for downstream applications (Fig 4). Additionally, we have developed a high throughput 96-well protocol that can be used for large-scale experiments and in an automated facility (Fig 3de). This is a critical step in the development of *S. meliloti* as a robust host for genome engineering and will facilitate its use in synthetic biology applications.

**Fig 3.**
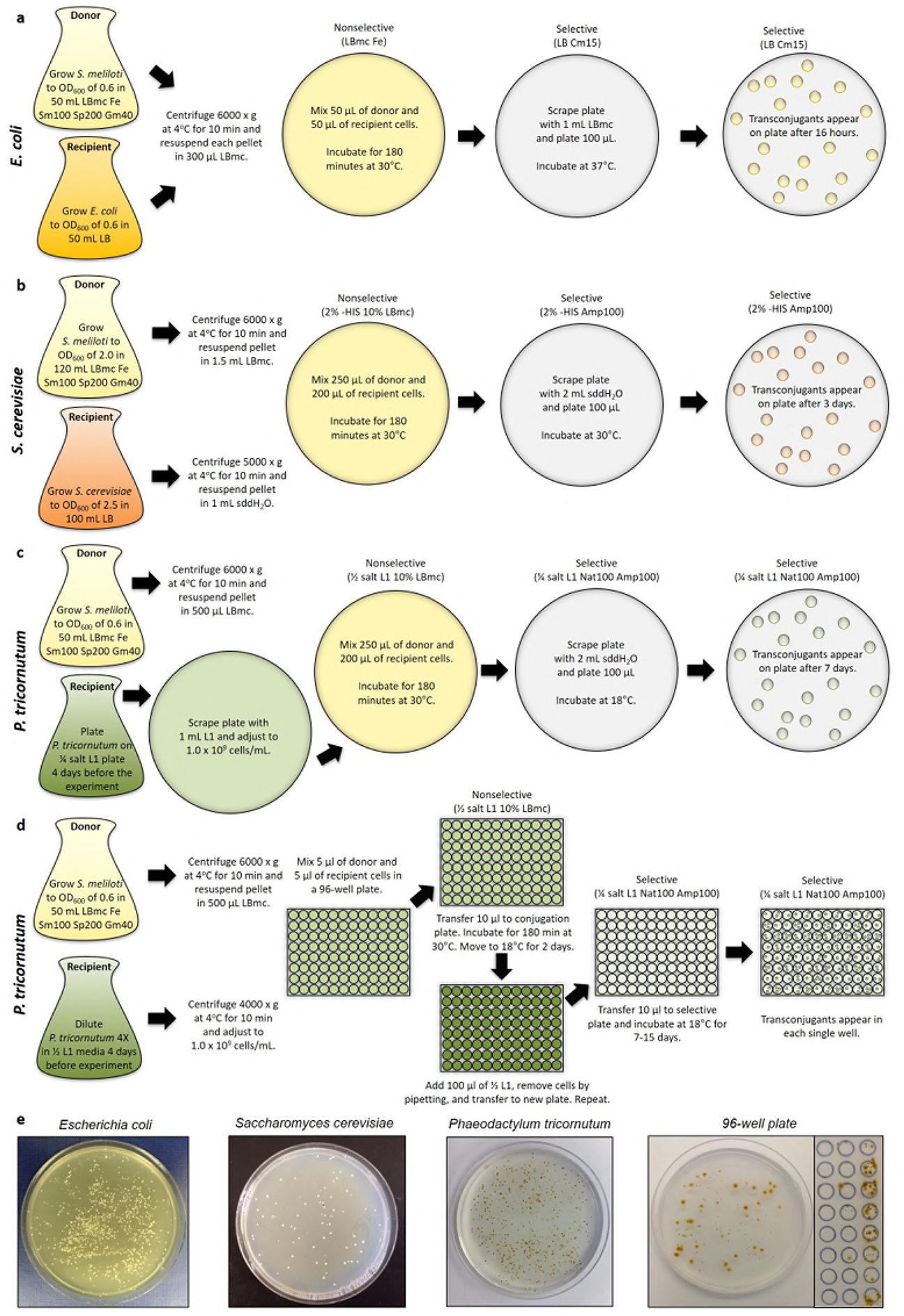
Optimized conjugation protocols from *S. meliloti* RmP4098 host strain to various recipient organisms. (a) Schematic of protocol for conjugation of pAGE1.0 from *S. meliloti* to *E. coli*. (b) Schematic of protocol for conjugation of pAGE1.0 plasmid from *S. meliloti* to *S. cerevisiae.* (c) Schematic of protocol for conjugation of pAGE1.0 plasmid from *S. meliloti* to *P. tricornutum* – standard protocol. (d) Schematic of protocol for conjugation of pAGE1.0 plasmid from *S. meliloti* to *P. tricornutum* – 96-well plate protocol. (e) Examples of plates containing final colonies that result from conjugation from *S. meliloti* to *E. coli, S. cerevisiae*, and *P. tricornutum* (standard or in a 96-well plate).

**Fig 4.**
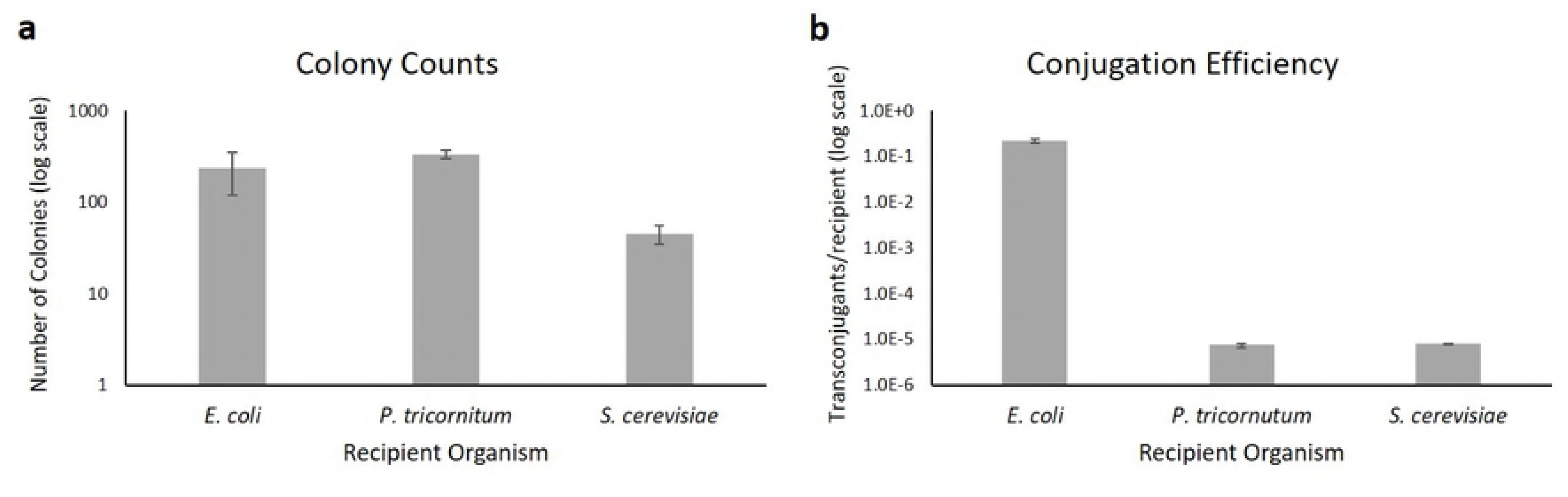
Summary of transconjugant counts and conjugation efficiency using *S. meliloti* as a donor to *E. coli, S. cerevisiae*, and *P. tricornutum.* (a) Number of transconjugants for each donor-recipient pair per experimental plate. The amount of conjugation mixture and ratio or donor to recipient per plate varies based on the different conjugation protocols. (b) Conjugation efficiency for each donor-recipient pair. Post-conjugation non-selective plates and pAGE selective plates were used to determine the conjugation efficiency (recorded as transconjugants/recipient) between each donor and recipient pair. Data is shown as bar plots with three biological and three technical replicates with error bars illustrating standard error of the mean.

Next, we evaluated the pAGE1.0 plasmids for DNA rearrangements and potential mutations that could have been introduced during conjugation from *S. meliloti* to *E. coli, S. cerevisiae* and *P. tricornutum*. In the first case, *E. coli* transconjugant plasmids were induced with 0.1% arabinose to obtain high copy number and directly isolated from *E. coli*. Plasmids from *S. cerevisiae* and *P. tricornutum* transconjugants were isolated and transformed into *E. coli*, then induced and once again isolated to obtain high quality DNA (pAGE1.0 replicates as a low-copy plasmid in *S. cerevisiae* and in *P. tricornutum* and it cannot be induced with arabinose to obtain high copy number within these species). Sixty exconjugants from each donor-recipient pair were selected and transformed to *E. coli*. DNA isolated from 59/60 *S. cerevisiae* colonies and 58/60 *P. tricornutum* colonies produced *E. coli* colonies (S4 Table). Then, 20 *E. coli* transformants (each transformed with DNA from independent *S. cerevisiae* colonies) and 30 *E. coli* transformants (each transformed with DNA from independent *P. tricornutum* colonies) were selected, induced with 0.1% arabinose, and the plasmid DNA was isolated. Diagnostic digests were performed on these plasmids with EcoRV-HF, and the number of plasmids with the expected banding pattern for *E. coli* was 18/20, *S. cerevisiae* 19/20 and *P. tricornutum* 16/30. (S5-S7 Figs) (S4 Table). Therefore, we can observe that correct pAGE1.0 plasmids can be rescued when conjugated from *S. meliloti* 90% of time in *E. coli*, 93% of the time in *S. cerevisiae*, and 52% of the time in *P. tricornutum*.

### Cloning of large high G+C content DNA fragment

With our MHS plasmids generated and protocols for direct transfer from *S. meliloti* in place, we were able to demonstrate the ability for our MHS plasmids to maintain large, high G+C content DNA in *S. meliloti. D. radiodurans* is an extremophile with a sequenced genome (G+C content 67%) and has many interesting characteristics including resistance to ionizing radiation and enhanced DNA-damage repair mechanisms [26]. We cloned the large 62 kb plasmid (CP1) (G+C content 57%) native to *D. radiodurans* R1 into pAGE2.0 that was transformed into *E. coli* and *S. meliloti* (Fig 5a). Diagnostic multiplex PCR and diagnostic digests using ApaI and ScaI were performed on *E. coli* and *S. meliloti* extracted DNA to verify the capture of the 62 kb *D. radiodurans* CP1 plasmid (Fig 5bc).

**Fig 5.**
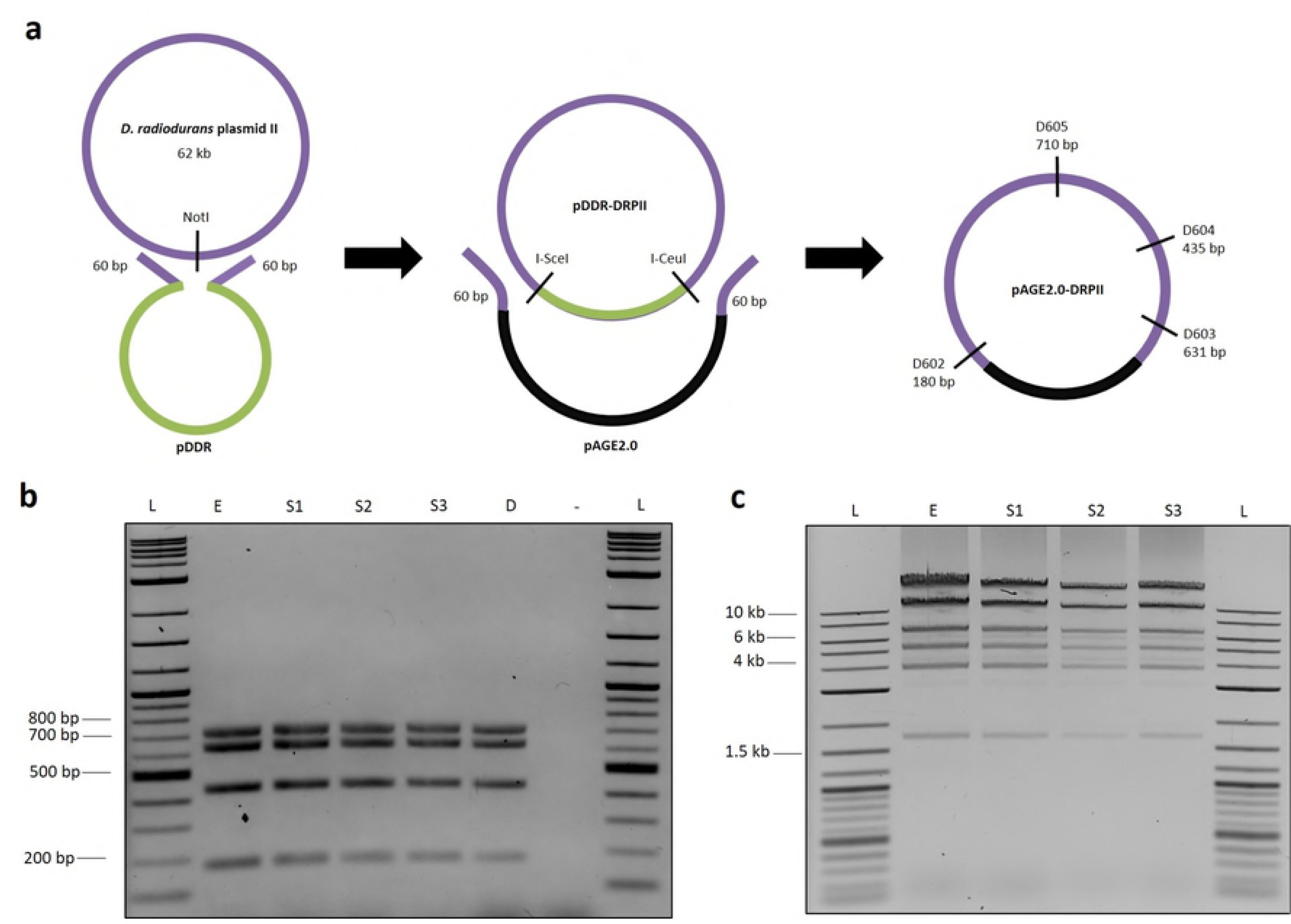
Cloning of 62 kb Plasmid II from *D. radiodurans* genome. (a) Schematic illustrating the strategy to clone the 62 kb *D. radiodurans* native plasmid into the pAGE2.0 plasmid. (b) Diagnostic MPX PCR of 62 kb *D. radiodurans* Plasmid II cloned into the pAGE2.0 plasmid after transformation into *E. coli* and *S. meliloti*. L, NEB 2-log ladder. E, pAGE2.0-DradPII extracted from *E. coli* Epi300. S1, S2, S3, pAGE2.0-DradPII extracted from *E. coli* after transformation from *S. meliloti* extracted DNA. D, *D. radiodurans* genomic DNA. -, no DNA negative control. (c) Diagnostic digest of 62 kb *D. radiodurans* Plasmid II cloned on the pAGE2.0 plasmid with ApaI and ScaI after transformation into *E. coli* and *S. meliloti*. L, NEB 2-log ladder. E, pAGE2.0-DradPII extracted from *E. coli* Epi300. S1, S2, S3, pAGE2.0-DradPII extracted from *E. coli* after transformation from DNA from three *S. meliloti* clones.

In summary, MHS plasmids were developed and shown to be stably maintained in *S. meliloti, E. coli, S. cerevisiae*, and *P. tricornutum*. These multi-host and multi-functional plasmids have promising applications for the cloning of large, high G+C content DNA fragments and their direct transfer to target organisms, as shown by the cloning of a 62 kb plasmid from *D. radiodurans*. As well, the pAGE/pBGE plasmids can be tailored for transfer to a target organism of interest, by including an origin of replication and selectable marker compatible with that organism. Conjugation from *S. meliloti* to *E. coli, S. cerevisiae*, and *P. tricornutum*, as well as showing *S. meliloti* as a conjugative recipient, illustrates that *S. meliloti* can be a suitable host organism for transfer of DNA with many model organisms, some of which are already used as host organisms. As well, 96-well plate conjugation from *S. meliloti* to *P. tricornutum* is promising for the future automation of these protocols. Due to the range of target organisms, it is possible that other target organisms can be added in the future, and the direct conjugative transfer from the *S. meliloti* host could contribute greatly to the field of constructing and installing synthetic chromosomes. These hosts will be invaluable for the cloning and installation of genes or large biosynthetic pathways, the study of organisms lacking genetic tools, and specifically for applications in industrially and agriculturally important strains (plants, marine organisms, soil microbiomes, etc.).

## Materials and Methods

### Microbial strains and growth conditions

*Sinorhizobium meliloti* strains was grown at 30°C in LBmc supplemented with appropriate antibiotics (streptomycin (100 μg mL^-1^), spectinomycin (200 μg mL^-1^), gentamicin (40 μg mL^-1^), tetracycline (5 μg mL^-1^), neomycin (100 μg mL^-1^), and 38 μM FeCl_3_ and/or 2 μM CoCl_2_, when appropriate.

*Saccharomyces cerevisiae* VL6-48 (ATCC MYA-3666: MATα his3-Δ200 trp1-Δ1 ura3-52 lys2 ade2-1 met14 cir^0^) was grown at 30°C in rich medium (2X YPD) or yeast synthetic complete medium lacking histidine supplemented with adenine (Teknova, Inc.) [27].

*Escherichia coli* (Epi300, Epicenter) was grown at 37°C in Luria Broth (LB) supplemented with appropriate antibiotics (chloramphenicol (15 μg mL^-1^)). *Escherichia coli* (ECGE101) was grown at 37°C in Luria Broth (LB) supplemented with appropriate antibiotics (chloramphenicol (15 μg mL^-1^), gentamicin (20 μg mL^-1^)), and diaminopimelic acid (60 μg mL^-1^).

*Phaeodactylum tricornutum* (Culture Collection of Algae and Protozoa CCAP 1055/1) was grown in L1 medium without silica at 18 °C under cool white fluorescent lights (75 μE m^-2^ s^-1^) and a photoperiod of 16 h light:8 h dark. Media was prepared as previously described [28].

*Deinococcus radiodurans* (R1) was grown at 30°C in TGY media.

### Development of Δ*hsdR S. meliloti* strains

Designer *S. meliloti* strains used in this study were created by taking the strain RmP3500, which lacks pSymA and pSymB where the *engA-tRNA^Arg^-rmlC* genomic region has been introduced into the chromosome [29] and re-introducing either pSymA or pSymB or both. Specifically, RmP4098, was made by re-introducing pSymB from RmP3491 into the strain RmP3910, which is derived from RmP3500, to create the strain RmP3950. From there, a Nm resistance cassette from strain RmP3975 was transduced into strain RmP3950. Double homologous recombination of the Nm cassette from RmP3975 into the genome of RmP3950 was selected on Nm Sm plates. The resulting strain, RmP4098, is ΔpSymA pSymB+ *hsdR*::Nm, Nm^R^ Sm^R^.

RmP3975 is a strain where the *hsdR* restriction gene has been replaced by a Nm resistance cassette. This strain was created by first PCR amplifying a downstream region and upstream region of the *hsdR* gene and cloning these PCR products into the *XbaI* site of pUCP30T using SLiC [30], then transforming into chemically competent DH5α. This construct was then verified via sequencing using M13 universal primers. The pUCP30T plasmid (GmR) with the *hsdR* upstream/downstream regions was then transformed into an *E. coli* strain harboring pKD46 (AmpR) which includes genes for lambda red recombinase, the resulting strain was named M2453. A Km/Nm cassette for *hsdR* deletion was then PCR amplified using high-fidelity DNA polymerase and purified. The cassette consisted of the antibiotic resistance gene flanked by regions of homology to the *hsdR* regions cloned into pUCP30T. The cassette was then electroporated into competent M2453 cells (grown in 1mM Arabinose to induce lambda red recombinase genes) and selected using Km (25 μg mL^-1^). Cells were then patched on Km (25 μg mL^-1^), Gm (10 μg mL^-1^) and Amp (100 μg mL^-1^). A KmR, GmR, and AmpS strain was then streak purified and named M2459. The resulting plasmid was then conjugated into RmP110 as recipient and M2459 as donor. Selection was done in Sm (200 μg mL^-1^) Nm (200 μg mL^-1^) plates. Resulting transconjugants were then patched on Gm (10 μg mL^-1^), (Nm 200 μg mL^-1^) and Sm (200 μg mL^-1^) plates and a Gm sensitive colony was streak purified (indicating double recombination of cassette with *hsdR* locus). The strain was then verified using diagnostic primers that spanned the *hsdR* upstream/downstream region with the Km/Nm cassette.

The Nm resistance cassette was then able to be transduced into the *S. meliloti* strains containing various combinations of pSymA and pSymB and selecting for Nm resistance. The Nm resistance cassette was then excised by introducing Flp recombinase to flip out the cassette using flanking FRT sites. The resulting strains are are RmP4258 (pSymA- pSymB- Δ*hsdR*), RmP4260 (pSymA+ pSymB- Δ*hsdR*), RmP4124 (pSymA- pSymB+ *Δ*hsdR), and RmP4125 (pSymA+ pSymB+ Δ*hsdR*).

### Development of *E. coli* ECGE101 Δ*dapA* strain

The *dapA* gene replacement with lambda pir and an erythromycin cassette from strain βDH10B was replicated in Epi300, resulting in a strain containing *trfA* and requirement of diaminopimelic acid (DAP) supplementation for growth. This is useful because it allows for replication of plasmids with RK2 oriV and makes it a convenient donor in biparental conjugation because the DAP requirement eliminates the need for antibiotic counter-selection. A lambda red recombinase plasmid (pKD46) was electroporated into Epi300, because Epi300 is recA-. The *dapA* region from βDH10B was amplified using the flanking primers DAP1 and DAP2. The fragment was electroporated into *E. coli* Epi300 containing pKD46 and transformants were selected on LB with DAP (60 μg mL^-1^) and erythromycin (200 μg mL^-1^), and inability to grow in the absence of DAP was confirmed. Such transformants were cured of pKD46 by growing at 37°C and confirming ampicillin (100 μg mL^-1^) sensitivity.

### Plasmid construction (pAGE/pBGE multi-host shuttle plasmids)

Briefly, pAGE and pBGE plasmids were constructed based on the pCC1BAC vector to which elements for replication in yeast were added. This backbone was amplified from plasmid p0521s (created at Venter institute and available on Addgene). This will allow replication and selection in yeast and *E. coli.* Additionally, it is low copy plasmid that can be induced to high copy with arabinose. Other components amplified for plasmid assembly include the antibiotic resistance cassettes for selection in *S. meliloti, repA2B2C2* or *repA1B1C1* for replication in *S. meliloti*, origin of transfer *(oriT)* for conjugation, and selective marker for algae (Ntc). The six fragments were PCR amplified and assembled in yeast using a yeast spheroplast transformation method. Next, DNA was isolated from yeast and moved to *E. coli* strain PG5alpha and genotyped using a multiplex PCR screen. Correct plasmids were moved from PG5alpha to Epi300 and Epi300 pTA-Mob. Diagnostic digests were performed to confirm correct assembly of plasmids.

### Plasmid stability assay of pAGE2.0

To characterize the ability of the Δ*AhsdR* strains to maintain these large pAGE plasmids, stability assays were performed to determine how long pAGE2.0 can be maintained in RmP4122 ΔpSymAΔ*hsdR*Nm^s^. This was performed in triplicate. Single colonies harboring pAGE2.0 were inoculated into LBmc FeCl_3_ + CoCl_2_ with Tc (5μg mL^-1^) at 30°C. The next day, 100uL of culture diluted to 10^-6^ was plated on LBmc FeCl_3_ + CoCl_2_ and grown 3 days at 30°C Cultures were subcultured 1000X in LBmc FeCl_3_ + CoCl_2_ without antibiotics and grown overnight. Approximately 10 doublings occurred per day. The next day, cultures were diluted to 10^-6^ and again 100μL was plated on LBmc FeCl_3_ + CoCl_2_. Cultures were subcultured again as before. When visible, 100 colonies were patched onto LBmc FeCl_3_ + CoCl_2_ with Tc (5μg mL^-1^) and without Tc as a control to ensure colony viability. These plates were incubated at 30°C for 3 days. The number of patched colonies that were able to grow on selective media was then recorded. The experiment was performed for 5 days to assess stability of pAGE2.0.

### Electroporation of *S. meliloti*

#### Preparation of competent *S. meliloti* cells for electroporation

Grow 500 mL of *S. meliloti* overnight in LBmc, 38 µM FeCl_3_, and streptomycin 100 μg mL^-1^ at 30 °C with shaking incubation to an OD_600_ of 2.0. Incubate flask on ice for 10 min then pellet at 6000 x g at 4°C for 10 min. Resuspend cells in 250 mL of sddH_2_O by gentile agitated in a water bath, top up volume to 500 mL, then pellet with the same conditions. Repeat this wash step with sddH_2_O two additional times, and then repeat once more with the same volume of 10% glycerol. Resuspend the pellet in 3 mL of 10% glycerol, flash freeze 200 µL aliquots and store at −80 °C.

#### Electroporation of *S. meliloti*

Incubate frozen cells on ice until fully thawed (about 15 min). Add 50 ng of DNA to 50 µL of competent cells in a 1.5 mL Eppendorf tube on ice, flick to mix, and incubate on ice for 5 min. Add transformation mixture to a 0.1 mm path length cuvette on ice and electroporate at 1.8 kV. Immediately add 1 mL LBmc 38µM FeCl_3_ and recover in a test tube at 30°C with shaking incubation (225rpm) for 120 min. Spread 500 µL of the transformation mixture on LBmc plates containing 38 µM FeCl_3_, 100 μg mL^-1^ streptomycin, and an appropriate concentration of antibiotic selection based on the transformed DNA. Incubate plates at 30°C for 3 days to allow for colony formation.

### PEG-mediated transformation of *S. meliloti*

#### Preparation of competent *S. meliloti* cells for PEG-mediated transformation

*S. meliloti* cells were cultured overnight in LBmc, 38µM FeCl_3_, and 100 μg mL^-1^ streptomycin at 30 °C with shaking incubation (225rpm). Upon reaching OD_600_ = 0.4, cells were harvested into sterile 500 mL centrifuge bottles, incubated on ice for 10 minutes, and pelleted at 4000 x g and 4°C for 10 minutes. Cells were then resuspended in 100 mL of ice-cold 100 mM CaCl_2_ by gentle pipetting and incubated on ice for an additional 30 minutes. Following incubation, cells were pelleted again at 4000 x g and 4°C for 10 minutes and resuspended in 1.25 mL of ice-cold 100 mM CaCl_2_ + 15% glycerol. The final resuspension volume was split into 25 µL aliquots, flash frozen using liquid nitrogen, and stored at −80°C for later use.

#### PEG-mediated transformation of *S. meliloti*

Frozen cells (25 µL aliquots) were incubated on ice until fully thawed. 200 ng of supercoiled pAGE DNA and 25 µL of 10% PEG 4000 were then added to the reaction tube and mixed evenly by gentle pipetting. Cells were then incubated on ice for 30 minutes, transferred to a 40°C water bath for 8 minutes, and immediately placed back on ice for 10 minutes. Following, 500 µL of LBmc, 38µM FeCl_3_ was added to the reaction tube and cells were recovered at 30°C with shaking incubation (225rpm) for 90 minutes. Following recovery, 250 µL of the transformation mixture were spread on LBmc plates containing 38 µM FeCl_3_, 100 μg mL^-1^ streptomycin, and an appropriate concentration of antibiotic selection. Plates were incubated at 30°C for 3 days to allow for colony formation.

### Transfer of DNA from *E. coli* to *S. meliloti* via conjugation

#### Preparation of *S. meliloti* (RmP4122) cells

An overnight culture (OD_600_=2.0) in LBmc, 38µM FeCl_3_, streptomycin 100 μg mL^-1^ was diluted 20x to make 20 mL culture and grown 6 hours shaking at 30°C in LBmc supplemented with streptomycin 100 µg mL^-1^ and Fe to OD_600_ of 0.9. The culture was diluted 2000X and grown with shaking at 30°C in 50 mL LBmc supplemented with streptomycin 100 µg mL^-1^ and Fe to OD_600_ of 0.6. The culture was centrifuged for 10 min at 6,000 x g at 4°C and resuspended in 300 µL of LBmc media.

#### Preparation *of E. coli* (ECGE 101 pTA-Mob pAGE1.0) cells

Saturated overnight culture of *E. coli* was diluted 20X into 50 mL LB supplemented with diaminopimelic acid 60 µg mL^-1^, chloramphenicol 15 µg mL^-1^, and gentamicin 20 µg mL^-1^ and grown with shaking at 37°C to OD_600_ of 0.6. The culture was centrifuged for 10 min at 6,000 x g at 4°C and resuspended in 300 µL of LBmc media.

#### Conjugation from *E. coli* to *S. meliloti*

50 µL of *E. coli* cells and 50 µL of *S. meliloti* cells were mixed directly on LBmc plates supplemented with 38µM FeCl_3_ and diaminopimelic acid 60 µg mL^-1^ and incubated for 180 minutes at 30°C. 1 mL of LBmc media was added to plates, cells were scraped, and 100 µL (from a dilution series of 10^-3^ to 10^-9^) was plated on LBmc plates supplemented with Fe, streptomycin 100 µg mL^-1^, and spectinomycin 200 µg mL^-1^ (Note: plates should be at least 35 mL thick). Plates were incubated at 30°C for 3 days before colonies are counted.

### Transfer of DNA from *S. meliloti* to *E. coli* via conjugation

#### Preparation of *S. meliloti* (RmP4098 pTA-Mob pAGE1.0) cells

Stock culture (OD_600_=2.0) was diluted 20X to make 20 mL culture and grown 6 hours shaking at 30°C in LBmc supplemented with streptomycin 100 µg mL^-1^ spectinomycin 200 µg mL^-1^, gentamicin 40 µg mL^-1^ and Fe to OD_600_ of 0.3. The culture was diluted 500X and grown with shaking at 30°C in 50 mL LBmc supplemented with streptomycin 100 µg mL^-1^, spectinomycin 200 µg mL^-1^, gentamicin 40 µg mL^-1^ to OD_600_ of 0.6. The culture was centrifuged for 10 min at 6,000 x g at 4°C and resuspended in 300 µL of LBmc media.

#### Preparation of *E. coli* (Epi300) cells

Saturated overnight culture of *E. coli* was diluted 20X into 50 mL LB and grown with shaking at 37°C to OD_600_ of 0.6. The culture was centrifuged for 10 min at 6,000 x g at 4°C and resuspended in 300 µL of LBmc media.

#### Conjugation from *S. meliloti* to *E. coli*

50 µL of *E. coli* cells and 50 µL of *S. meliloti* cells were mixed directly on LBmc plates supplemented with Fe and incubated for 180 minutes at 30°C. 1 mL of LBmc media was added to plates, cells were scraped, and 100 µL (from a dilution series of 10^-3^ to 10^-9^) was plated on LB plates supplemented with chloramphenicol 15 µg mL^-1^. Plates were incubated at 37°C for 16 hours before colonies are counted.

### Transfer of DNA from *S. meliloti* to *S. cerevisiae* via conjugation

#### Preparation of *S. meliloti* (RmP4098 pTA-Mob pAGE1.0) cells

Stock culture (OD_600_=2.0) was diluted 20x to make 20 mL culture and grown 6 hours shaking at 30°C in LBmc supplemented with streptomycin 100 µg mL^-1^, spectinomycin 200 µg mL^-1^, gentamicin 40 µg mL^-1^ and 38µM FeCl_3_ to OD_600_ of 0.3. The culture was diluted and grown with shaking at 30°C in 120 mL LBmc supplemented with streptomycin 100 µg mL^-1^, spectinomycin 200 µg mL^-1^, gentamicin 40 µg mL^-1^, acetosyringone 100 µg mL^-1^ to OD_600_ of 2.0 (add arabinose 100 µg mL^-1^ to the growing culture 1 hour before the target OD is reached). The culture was centrifuged for 10 min at 6,000 x g at 4°C and resuspended in 1.5 mL of LBmc media.

#### Preparation of *S. cerevisiae* (VL6-48) cells

100 mL of liquid grown culture was grown with shaking at 30°C in 2X YPAD media to OD_600_ of 2.5. The culture was centrifuged for 10 min at 5,000 x g and resuspended in 1 ml of H_2_0.

#### Conjugation from *S. meliloti* to *S. cerevisiae*

200 µL of *S. cerevisiae* cells and 250 µL of *S. meliloti* was directly mixed on a 2% -HIS plate supplemented with 10% LBmc, 38µM FeCl_3_ and acetosyringone 100 µg mL^-1^ (Note: plates were dried out in the hood for 1 hour prior to conjugation). Then plates were incubated for 180 minutes at 30°C. Then, 2 mL of sddH_2_0 was added to plates and cells were scraped. 100 µL of the scraped cells was plated on 2% -HIS plates supplemented with ampicillin 100 µg mL^-1^. Plates were incubated at 30°C where colonies start to appear after 2-3 and colonies are counted after 5 days.

### Transfer of DNA from *S. meliloti* to *P. tricornutum* via conjugation

#### Preparation of *P. tricornutum* cells

250 µL of liquid grown culture was adjusted to 1.0 x 10^8^ cells mL^-1^ using counts from a hemocytometer, was plated on ½L1 1% agar plates and grown for 4 days. 1 mL of L1 media was added to the plate and cells were scraped, counted using a hemocytometer, and adjusted to a concentration of 1 x 10^9^ cells mL^-1^.

#### Preparation of *S. meliloti* (strain A-R-, pAGE1.0, pTA-Mob) cells

Stock culture (OD_600_=2.0) was diluted 20x to make 20 mL culture and grown 6 hours shaking at 30°C in LBmc supplemented with spectinomycin 200 µg mL^-1^, gentamicin 40 µg mL^-1^ and Fe to OD_600_ of 0.3. The culture was diluted 25X and grown for 12 hours with shaking at 30°C in 50 mL LBmc supplemented with spectinomycin 200 µg mL^-1^, gentamicin 40 µg mL^-1^ to OD_600_ of 0.6. The culture was centrifuged for 10 min at 5,000 x g at 4°C and resuspended in 500 µL of LBmc media.

#### Conjugation from *S. meliloti* to *P. tricornutum*

200 µL of *P. tricornutum* cells and 200 µL of *S. meliloti* cells were mixed directly on ½L1 10% LBmc 1% agar plates (Note: plates are dried in the biosafety cabinet for one hour before conjugation) and incubated for 180 minutes at 30°C in the dark, then moved to 18°C in the light and grown for 2 days. After two days, 2 mL of L1 media was added to plates, cells were scraped, and 100 µL (5%) was plated on ¼L1 1% agar plates supplemented with nourseothricin 100 µg mL^-1^, and ampicillin 100 µg mL^-1^ (Note: plates should be at least 35 mL thick). Plates were incubated at 18°C in the light/dark cycle and colonies start to appear after 7 days and are allowed to develop to 14 days before colonies are counted.

### Transfer of DNA from *S. meliloti* to *P. tricornutum* via conjugation in a 96-well plate

#### Preparation of *P. tricornutum* cells

200 µL of liquid grown culture was diluted using counts from a hemocytometer, and grown in ½L1 media for 4 days. Cell counts from a hemocytometer were used and culture was pelleted at 4000 x g 10 min 4°C, and adjusted to a concentration of 1 x 10^9^ cells mL^-1^.

#### Preparation of *S. meliloti* (strain A-R-, pAGE1.0, pTA-Mob) cells

Stock culture (OD_600_=2.0) was diluted 20x to make 20 mL culture and grown 6 hours shaking at 30°C in LBmc supplemented with spectinomycin 200 µg mL^-1^, gentamicin 40 µg mL^-1^ and Fe to OD_600_ of 0.3. The culture was diluted 25X and grown for 12 hours with shaking at 30°C in 50 mL LBmc supplemented with spectinomycin 200 µg mL^-1^, gentamicin 40 µg mL^-1^ to OD_600_ of 0.6. The culture was centrifuged for 10 min at 5,000 x g at 4°C and resuspended in 500 µL of LBmc media.

#### Conjugation from *S. meliloti* to *P. tricornutum*

5 µL of *P. tricornutum* cells and 5 µL of *S. meliloti* cells were mixed together in a 96-well plate. The mixture (10 µL) was transferred to a 96-well plate containing 200 µL of ½L1 10% LBmc 1% agar (note: plates are dried in the biosafety cabinet for one hour before conjugation). This conjugation plate was incubated for 180 minutes at 30°C in the dark, then moved to 18°C in the light and grown for 2 days. After two days, 100 µL of L1 media was added to wells and cells were scraped (X2), and 10 µL (5%) was plated on ¼L1 1% agar supplemented with nourseothricin 100 µg mL^-1^, and ampicillin 100 µg mL^-1^ in a 96-well plate. Plates were incubated at 25°C in the light/dark cycle for 24 hours and then 18°C in the light/dark cycle for an additional 24 hours. Colonies start to appear after 7 days and are allowed to develop up to 14 days before colonies are counted.

#### Plasmid DNA isolation

Plasmid DNA (<60 kb) was isolated from *E. coli* using the BioBasic EZ-10 miniprep kit. Plasmid DNA was isolated from all other species using a modified alkaline lysis protocol. Plasmid DNA was isolated from all other species and *E. coli* containing plasmids >60 kb using the modified alkaline lysis protocol described below. Steps 1–3 are variable depending on the species, while steps 4–10 are common for all species. Steps 1–3 for *E. coli* and *S. meliloti*. (1) Five mL cultures were grown to saturation overnight. (2) Cells were pelleted at 5000 x g for 10 min at 4°C, and the supernatant was discarded. (3) Cells were resuspended in 250 uL of resuspension buffer (which contained 240 ml P1 (Qiagen), 5 ml of 1.4 M b-Mercaptoethanol and 5 ml Zymolyase solution (Zymolyase solution: 200 mg Zymolyase 20 T (USB), 9 ml H2O, 1 ml 1 M Tris pH7.5, 10 ml 50% glycerol, stored at 20 °C). Steps 1–3 for *S. cerevisiae*. (1) Five mL of culture was grown to saturation. (2) Cells were pelleted at 5000 x g for 10 min 10 min at 4°C, and the supernatant was discarded. (3) Cells were resuspended in 250 mL resuspension buffer (as described above) and incubated at 37 °C for 60 min. Steps 1–3 for *P. tricornutum*. (1) Five mL cultures were harvested during exponential growth phase. (2) Cells were pelleted at 4,000g for 10 min at 4°C, and the supernatant was discarded. (3) Cells were resuspended in 250 mL resuspension buffer, which contained 235 ml P1 (Qiagen), 5 ml hemicellulase 100 mg ml 1, 5 ml of lysozyme 25 mg ml 1, and 5 ml Zymolyase solution (Zymolyase solution: 200 mg Zymolyase 20T (USB), 9 ml H2O, 1 ml 1 M Tris pH7.5, 10 ml 50% glycerol, stored at 20 °C) and then cells were incubated at 37 °C for 30 min. Steps 4–10 common for all species. (4) 250 uL of lysis buffer P2 (Qiagen) was added and samples were inverted 5-10 times to mix. (5) 250 ml of neutralization buffer P3 was added and samples were inverted 5-10 times to mix. (6) Then samples were spun down at 16,000g, 10 min at 4°C (7) Supernatant was transferred to a clean tube and 750 ul ice-cold isopropanol was added and the samples were mixed by inversion and spun down at 16,000g, 10 min at 4°C (8) Next the supernatant was removed and 750 ul ice-cold 70% EtOH was added and samples were mixed by inversion and spun down at 16,000g, 5 min. (9) Next the supernatant was discarded, pellets were briefly dried and resuspended in 50 uL of TE buffer. (10) After that the samples were kept at 37 °C for 30-60 min to dissolve.

#### Agarose plug DNA isolation

DNA from *Deinococcus radiodurans* for Transformation-Associated Recombination (TAR) cloning [23] of the 62-kb plasmid was isolated in agarose plugs using the Bio-Rad CHEF Genomic DNA Plug Kit with an adapted protocol. To prepare the plugs, 50 mL *of D. radiodurans* culture was grown to OD_600_ of 1.0, chloramphenicol (100 μg mL^-1^) was added and the culture was grown for an additional hour. The culture was centrifuged at 5000 × g for 5 min at 10°C. Cells were washed once with 1 M sorbitol in 1.5 mL Eppendorf tubes and centrifuged at 4000 RPM for 3 min. The supernatant was removed. Cells were resuspended in 600 uL of protoplasting solution. The cell suspension was incubated for 5 min at 37°C and mixed with an equal volume of 2.0% low-melting-point agarose in 1 × TAE buffer (40 mM Tris, 20 mM acetic acid and 1 mM EDTA) which was equilibrated at 50°C. Aliquots of 95 μl were transferred into plug molds (Bio-Rad, catalog # 170-3713) and allowed to solidify for 10 min at 4°C. Next, plugs were removed from the molds into 50 ml conical tube containing 5 mL of protoplasting solution (for 10 mL: 4.56 mL of SPEM solution, 1000 μl Zymolyase-20 T solution (50 mg mL^-1^ dissolved in H_2_O), 400 μl lysozyme (25 mg ml^-1^), 400 μl Hemicellulase (25 mg mL^-1^), 50 μl β-Mercaptoethanol) and incubated for 45 min at 37°C. Next, plugs were washed with 25 ml of wash buffer (20 mM Tris, 50 mM EDTA, pH 8.0), and then incubated in 5 ml in Proteinase K buffer (100 mM EDTA (pH 8.0), 0.2% sodium deoxycholate, and 1% sodium lauryl sarcosine, 1 mg ml^-1^ Proteinase K) for 24 hr at 50°C. Wash the plugs 4 times with 25 mL of wash buffer (20 mM Tris, pH 8.0, 50 mM EDTA) for 30 minutes each at room temperature. Leave it in wash buffer overnight. The next day, transfer 1-2 plugs to a 1.5 mL Eppendorf tube and wash it 4 times with 10X diluted wash buffer for 30 minutes each. Store at 4°C in 1.5 mL tube of diluted wash buffer. To isolate the DNA from the plug, wash once with 10X diluted wash buffer for 1 hour. Wash once with TE buffer for 1 hour, then remove all the TE. Put the 1.5 mL tube with the plug in a 42°C water bath for 10 mins. Then transfer to 65°C water bath for 10 mins. Return to 42°C water bath, wait for 5-10 mins then add 50 μl of TE buffer followed by 3 μl of agarose and leave it at 42°C for an hour. Add 50 μl TE buffer and leave it for 2-4 hours. Run 1-2 μl of this on 1% gel.

#### Cloning 62 kb CP1 plasmid from *Deinococcus radiodurans*

High quality total genomic DNA from *D. radiodurans* was isolated using low melting point agar plugs. To clone the CP1 plasmid from *D. radiodurans*, a single cut restriction enzyme (NotI) site was identified and used to linearize the plasmid. A 200 bp sequence on either side of the NotI cut site was amplified and inserted into Designer Microbes Inc proprietary plasmid using yeast assembly, to create regions of homology to CP1. The resulting plasmid was called pDRR. Next, TAR cloning [23] was performed using the linearized plasmids with homology to CP1 and *D. radiodurans* genomic DNA digested with NotI, resulting in yeast clone pDDR c1. Sub-cloning of CP1 was carried through yeast assembly using the linearized pAGE2.0 with hooks, short homologous sequences, to *D. radiodurans* CP1 (pAGE2.0-D. rad-PII-HOOKS), along with I-CeuI and I-SceI digested product of pDDR c1. The resulting yeast colonies were pooled and DNA was extracted by modified alkaline lysis and transformed into *E. coli*. Following induction with arabinose, DNA was extracted from *E. coli* by alkaline lysis and subsequently electroporated into *S. meliloti*.

## Acknowledgements

This work was supported by a Genome Canada Grant (OGI-111) to T.C.C., T.M.F. and B.J.K. Research in the labs of D.R.E., T.C.C., T.M.F. and B.J.K. is also supported by Natural Sciences and Engineering Research Council of Canada (NSERC), RGPIN-2015-04800, RGPIN-2018-04754, RGPIN-2018-06481, RGPIN-2018-06172 respectively.

We would like to thank Dr. Murray Junop, Western University, London, On, Canada, for providing *Deinococcus radiodurans* (R1) strain.

Conflict of interest statement. B.J.K. is Chief Executive Officer of Designer Microbes Inc. and holds Designer Microbes Inc. stock.

## Supporting Information

**S1 Fig. Verification of *hsdR* deletion in *S. meliloti* by diagnostic PCR.** (a) Schematic of primer set locations at the *hsdR* gene locus in wildtype *S. meliloti*, when the *hsdR* gene is present, and when the *hsdR* gene is replaced with FRT-Km/Nm-FRT cassette, which was subsequently excised via Flp recombinase. The pink arrow indicates an FRT site following loss of the FRT-Km/Nm-FRT cassette. (b) Gel electrophoresis of diagnostic colony PCR conducted with primer sets A, B, and C on wildtype and designer *S. meliloti* strains. Expected band size for primer set A is 953 bp if the *hsdR* gene is present or absent. Expected band size for primer set B is 912 bp if the *hsdR* gene is present, and no band is expected if the *hsdR* gene is absent. Expected band size for primer set C is 4013 bp if the *hsdR* gene is present, and 787 bp if the *hsdR* gene is absent. L, 1 kb ladder. 1, RmP3909 ΔpSymAB Δ*hsdR*. 2, RmP3952 ΔpSymB Δ*hsdR*. 3, RmP3953 ΔpSymA Δ*hsdR*. 4, RmP3954 Δ*hsdR*. 5, Rm5000 ΔpSymA Δ*hsdR* Rif^R^. 6, RmP110 wildtype. 7, RmP110 wildtype isolated gDNA. 8, no DNA control.

**S2 Fig. Diagnostic digest of MHS plasmids.** Following yeast assembly, transformation *to E. coli* Epi300 cells, induction and plasmid DNA extraction, MHS plasmids (pAGE1.0, pAGE2.0, pAGE3.0, pBGE1.0, pBGE2.0, pBGE3.0) were digested with I-CeuI, I-SceI, PacI, and PmeI,

**S3 Fig. Plasmid stability assay of pAGE2.0 in *S. meliloti* over 50 generations.** Graph depicting the percentage of RmP4122 ΔpSymA Δ*hsdR* colonies, from three independent cultures originally containing pAGE2.0, unable to grow on selective media (LBmc 38 μM FeCl_3_ Tc 5 μg mL^-1^) after each subculturing event in non-selective media, to a total of approximately 50 generations.

**S4 Fig. PEG-mediated transformation of MHS plasmids into *S. meliloti*.** Experimental and control plates from PEG-mediated transformation of pAGE1.0, pAGE2.0 and pAGE3.0 into *S. meliloti* RmP4122 ΔpSymA *ΔhsdR.* pAGE1.0 transformation results were discarded due to the comparable number of colonies consistently observed on experimental and control plates.

**S5 Fig. EcoRV-HF diagnostic digest of pAGE1.0 plasmids extracted from 20 *E. coli* colonies following conjugation from *S. meliloti* to *E. coli.*** Expected band sizes are 10,288 bp, 5235 bp, 2377 bp, and 229 bp. L, 1 kb ladder. 1-20, the 20 pAGE1.0 plasmid extracts from *E. coli.*

**S6 Fig EcoRV-HF diagnostic digest of pAGE1.0 plasmids extracted from 20 *E. coli* colonies following conjugation from *S. meliloti* to *P. tricornutum*, plasmid isolation, transformation to *E. coli*, and plasmid induction in *E. coli.*** Expected band sizes are 10,288 bp, 5235 bp, 2377 bp, and 229 bp. L, 1 kb ladder. 1-20, the 20 pAGE1.0 plasmid extracts from *E. coli.*

**S7 Fig EcoRV-HF diagnostic digest of pAGE1.0 plasmids extracted from 20 *E. coli* colonies following conjugation from *S. meliloti* to *S. cerevisiae*, plasmid isolation, transformation to *E. coli*, and plasmid induction in *E. coli.*** Expected band sizes are 10,288 bp, 5235 bp, 2377 bp, and 229 bp. L, 1 kb ladder. 1-20, the 20 pAGE1.0 plasmid extracts from *E. coli.*

**S1 Table. Plasmid stability of pAGE2.0 in *S. meliloti* RmP4122 ΔpSymA Δ*hsdR*.** Determined by the number of colonies unable to grow on selective media (LBmc 38 μM FeCl_3_ Tc 5 μg mL^-1^) following subculturing in nonselective media (LBmc 38 μM FeCl_3_).

**S2 Table. Summary of DNA transfer to *S. meliloti* RmP4122 ΔA ΔR including PEG-mediated transformation, electroporation and conjugation of pAGE1.0, pAGE2.0 and pAGE3.0, where applicable.** Transformation efficiency for PEG-mediated and electroporation is reported as CFU μg^-1^ of DNA, conjugation efficiency is reported as transconjugants/recipient. Mean is the average of three biological and three technical replicates. *pAGE1.0 for the PEG-mediated transformation method consistently results in a similar number of colonies observed on the experimental and negative control plates (which can be seen in S4 Fig).

**S3 Table. Ratio of donor to recipient cells and conjugation efficiency (as transconjugants/recipient) for each donor and recipient pair.** Pre-conjugation plates selecting for the conjugative participants and colony counts were used to determine the ratio of donor to recipient organism going into the conjugation mixture.

**S4 Table. Analysis of pAGE1.0 plasmids recovered from conjugations from *S. meliloti* to *E. coli, S. cerevisiae*, and *P. tricornutum*.** Diagnostic digests of pAGE1.0 plasmids with EcoRV-HF, following transformation and induction in *E. coli*, where applicable.

**S5 Table. List of oligonucleotides used in this study.**

**S6 Table. List of strains used in this study.**

**S7 Table. List of plasmids used in this study.**

**Author Contributions**
T.C.C., T.M.F and B.J.K. conceived the project and assisted in design. S.L.B., M.M., T.H., R.C., R.M., M.Z., O.M., P.J., T.M.F and B.J.K. performed the experiments. S.L.B., P.J., D.R.E., T.C.C., T.M.F and B.J.K. conducted data analysis and interpretation. S.L.B., D.R.E., T.C.C., T.M.F. and B.J.K. wrote the manuscript.

